# Fine needle aspiration of human lymph nodes reveals cell populations and soluble interactors pivotal to immunological priming

**DOI:** 10.1101/2023.10.18.562983

**Authors:** Nicholas M. Provine, Adam Al-Diwani, Devika Agarwal, Kyla Dooley, Amelia Heslington, Andrew G. Murchison, Lucy C. Garner, Fintan Sheerin, Paul Klenerman, Sarosh R. Irani

## Abstract

Lymph node (LN) fine needle aspiration (LN FNA) represents a powerful technique for minimally invasive sampling of human lymph nodes *in vivo* and has been used to good effect to directly study aspects of the human germinal center response. However, systematic deep phenotyping of the cellular populations and cell-free proteins recovered by LN FNA has not been performed. Thus, we studied human cervical LN FNAs as a proof-of-concept and used single-cell RNA-sequencing and proteomic analysis to benchmark this compartment, define the purity of LN FNA material, and facilitate future studies into this immunologically pivotal environment. Our data provide evidence that LN FNAs contain *bone fide* LN-resident innate immune populations, with minimal contamination of cells or proteins from blood. Examination of these populations reveals unique biology not predictable from equivalent blood-derived populations. LN FNA supernatants represent a specific source of lymph- and lymph node-derived proteins, and can, in combination with transcriptomic approaches, identify likely receptor-ligand interactions. This study provides the first description of the types and abundance of immune cell populations and cell-free proteins that can be efficiently studied by LN FNA. These findings are of broad utility for understanding LN physiology both in health and disease, including infectious or autoimmune perturbations, and in the case of cervical nodes, neuroscience.

## INTRODUCTION

The first step in the induction of adaptive cellular and humoral immune responses requires the priming of naïve T and B cells. To increase the likelihood of interaction between rare naïve lymphocytes and their cognate antigen, and to localize critical accessory cell populations, this process primarily occurs within highly specialized secondary lymphoid organs (SLOs) – the spleen and peripheral lymph nodes (LNs). Priming often begins with the trafficking of migratory dendritic cells (DCs), carrying tissue antigens, through lymphatics to the LN (1). Thereafter, conventional DCs (cDCs) may process and present antigen in complex with HLA molecules to prime naïve T cells (2). Additionally, specialized LN stromal cells - follicular dendritic cells (FDCs) - capture and retain antigen-rich immune complexes for induction of B cell responses, which supports the development of the germinal center (GC) reaction with consequent B cell affinity maturation (3). Local soluble factors, including specific chemokines and cytokines, have a critical role in the priming of both T and B cell responses. Thus, the highly tissue-restricted nature of these processes means direct access to lymphoid tissue is a necessity to study immune priming events.

Several practical issues exist around sampling secondary lymphoid tissue in humans. Firstly, due to the anatomy of the lymphatics system, one, or a few, region-specific “draining” LNs may be specifically involved in any single immune priming process. For example, elegant work in recent years has established that cervical LNs of the head and neck drain cerebral lymphatics (4). Yet, it is not known how many of the seven distinct cervical LN basins are relevant to CNS immunology. Secondly, immune priming is a highly temporally regulated response and examination of lymphoid tissue in the steady state may bear little relation to the processes occurring during the early days of an immune response (5). Thus, to study disease-relevant events both the precise target LN and the time of sampling must be established.

To overcome these issues, in the past five years, ultrasound-guided lymph node fine needle aspiration (LN FNA) has been employed to allow minimally invasive, directly visualized, and time-resolved sampling of specific LNs in volunteers (6–14). This is particularly pertinent when mapping the spatial and time-dependent priming processes which occur in the GC response. We have previously applied LN FNA to ask whether cervical LNs are involved in the pathogenesis of autoantibody-mediated brain disease (6, 7). Others have utilized LN FNA in healthy volunteers to examine the GC response to HIV (8), influenza (10), or SARS-CoV-2 virus vaccines (9, 12–14). This approach has permitted studying both B cells and T follicular helper (T_FH_) cells, which support GC formation and maintenance (15). These studies all obtained robust lymphocyte populations and have focused on the effector phase of lymphocyte function within the LN. Innate immune cell populations, which are pertinent to both GC reactions and T cell priming, were not examined in detail.

Thus, it remains to be established if LN FNA is a suitable technique to examine the innate and adaptive immune cell interactions so critical for immune priming. Furthermore, with the exception of our focused investigations of CXCL13 (6) and autoantibody levels (7), in the context of immunology, LN FNA studies have primarily examined cellular fractions. Hence, it remains unclear if the technique is broadly suitable for studying soluble factors within LNs, including cytokines and antigens.

To address these questions, we performed single-cell RNA-sequencing (scRNA-seq) with surface protein quantification and high-dimensionality proteomic analysis of cervical LN FNA material and matched blood across four healthy volunteers. We found LN-resident innate immune populations can be efficiently recovered by FNA, and examination of these populations revealed plausible, LN-specific cell-cell interactions. Furthermore, proteomic analysis of cell-free LN FNA material identified proteins not detectable in blood, including candidate lymph-derived material, with utility for neuroimmunology studies. This study demonstrates the utility of LN FNA in addressing these aspects of immune priming directly in vivo using human volunteers and creates an early LN atlas of cells and soluble interactors with significant utility for multiple applications in human autoimmune diseases, cancers, and vaccinations.

## RESULTS

### Annotation of immune populations recovered from LN FNA

To investigate the feasibility of using LN FNA to study innate immune crosstalk we utilized single-cell RNA-sequencing (scRNA-seq) and high-dimensionality proteomics on cervical LN aspirates and matched peripheral blood from four healthy volunteers (Figure 1A). Our sorting strategy was devised to enrich for innate immune populations and removed the high proportion of naïve T and B cells present in the LN aspirates of our young, healthy participants. Three populations of cells were separately sorted from each sample: non-T/non-B (live, CD45^+^CD3^−^CD19^−^), non-naïve T cells (live, CD45^+^CD3^+^ and NOT CD45RA^+^CD27^+^), and non-naïve B cells (live, CD45^+^CD19^+^ and NOT IgD^+^CD27^−^) (Supplemental Figure 1). The three sorted populations were recombined at an approximately 1:1:1 ratio and libraries were prepared using the 10x Genomics platform.

**Figure 1.**
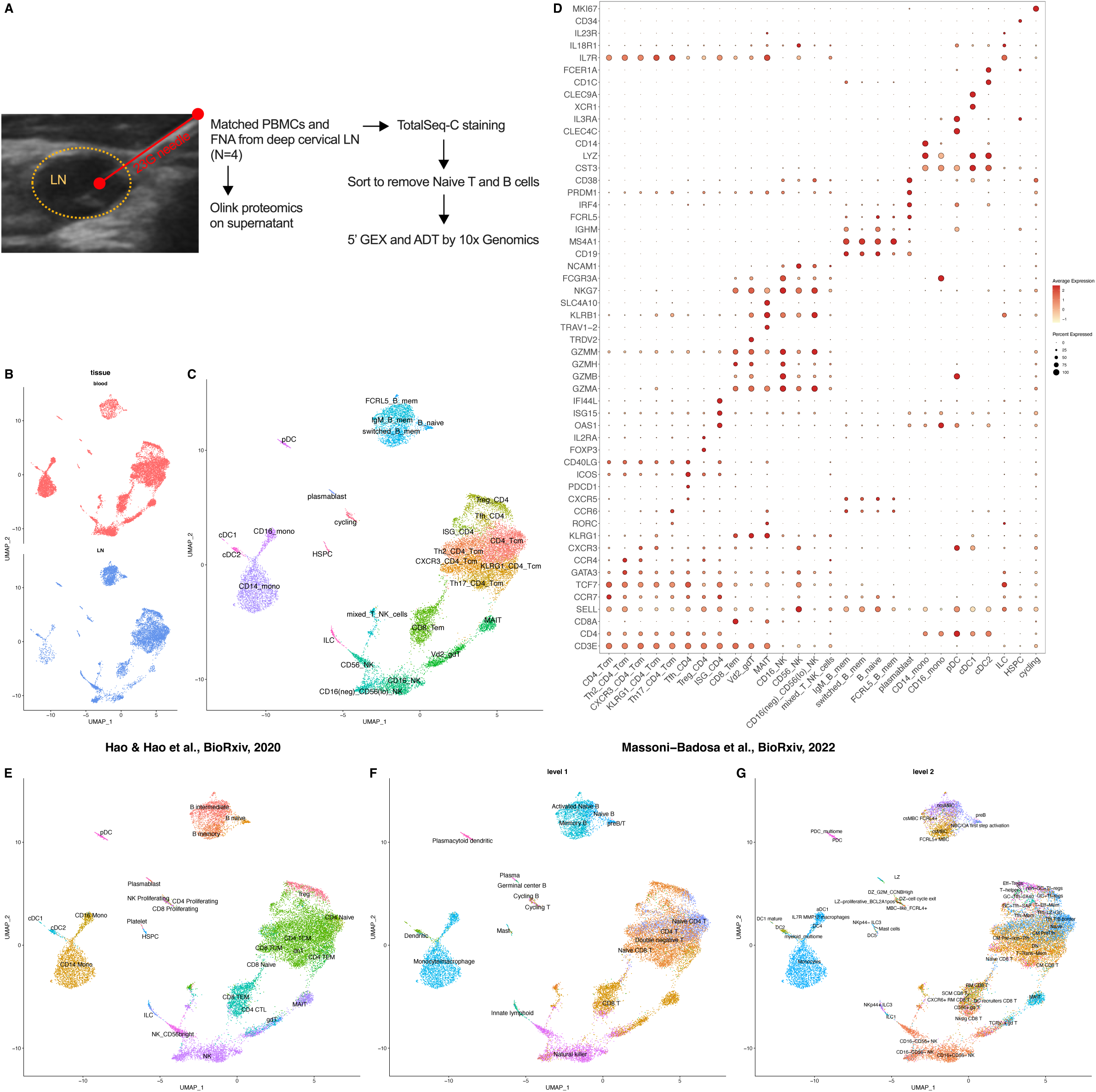
Identification of cell populations harvested by lymph node fine needle aspiration. **(A)** Experimental plan. Cervical LN material and matched peripheral blood was collected from *n* = 4 healthy donors. **(B)** UMAP of all cells (*n* = 19,154) separated by tissue of origin. **(C)** UMAP of all cells with cluster annotations. **(D)** Dot plot highlighting key markers used to define each cluster. **(E-G)** UMAP showing the annotation of populations based on the following reference datasets: (E) human PBMCs (16), (F) broad level 1 annotation of human tonsil, and (G) fine level annotation of human tonsil (17). The Azimuth package was used for reference mapping (see methods).

After data QC and clean-up, we had 19,154 high-quality cells (11,694 from PBMC and 7,460 from LN). Following clustering and batch correction, both tissue sources were found to contribute to all clusters (Figure 1B, 1C), although to varying degrees (discussed below). To annotate clusters, we utilized the Azimuth algorithm (16) to perform automated reference mapping using both a reference PBMC dataset (16), and a recently published atlas of human tonsil tissue (17). Annotations were refined using expert knowledge to give a total of 28 annotated clusters (Figure 1C and 1D), with ADT protein data used to assist cluster annotation (Supplemental Figure 2A).

Cluster annotation using the reference PBMC dataset L2 annotation (Figure 1E) generated high confidence scores for blood and LN-origin cells (Supplemental Figure 2B, 2C). Level 1 annotation of the reference tonsil dataset also gave high confidence mapping (Supplemental Figure 2B and C), but the annotations revealed limited granularity (Figure 1F). By contrast, reference mapping using the level 2 tonsil annotations identified multiple innate subsets in our samples which specifically associated with lymphoid tissue (Figure 1G). These were identified within dendritic cells (DCs), innate lymphocytes (ILCs) and natural killer (NK) cells. High resolution of the B cell and T cell compartments was also achieved, including identification of GC-associated B and T cell populations. However, given the 137 unique cluster annotations these mapped with lower confidence (Supplemental Figure 2B and 2C). Nonetheless, our data indicate an efficient recovery of multiple important innate immune populations within the LN.

To avoid over-fitting, we annotated clusters without over-reliance on the level 2 tonsil reference annotations. Consistent with the FACS analysis, no cells classifiable as naïve CD4^+^ or CD8^+^ T cells were identified, alongside only a small residual naïve B cell cluster (Figure 1C). We identified a T_FH_ cluster that uniquely expressed *PDCD1* (and CD279 protein) and *CXCR5*, and was highest for *ICOS* and *CD40LG* expression (Figure 1C and 1D, Supplemental Figure 2A). We clearly identified multiple innate immune populations: cDC1s (gene: *CST3, XCR1, CLEC9A;* protein: CD141^hi^), cDC2s (gene: *CST3, CD1C, FCER1A*; protein: CD1c^hi^), and plasmacytoid DCs (pDCs; gene: *CST3, CLEC4C, IL3R*; protein: CD303^hi^, CD123^hi^) (Figure 1C and 1D, Supplemental Figure 2A). Separate ILC (gene: *KLRB1, IL7R;* protein: Lineage^−^CD161^+^CD127^+^) and hematopoietic stem and progenitor cell populations were also observed (HSPCs; gene: *CD34*; protein: Lineage^−^) (Figure 1C and 1D, Supplemental Figure 2A). Within NK cells, we identified a population that expressed low levels of CD56 and CD16 (at gene and protein level), but still expressed other NK cell markers (e.g., *NKG7*), a phenotype previously associated with lymphoid tissue (17). Collectively, these data demonstrate recovery of innate immune cell populations from the LN in addition to other biologically relevant tissue-specific populations.

### Relative abundance of cell populations in blood versus lymph node

We noted that after more than one needle pass, aspirates could be discolored red, which was associated with a red cell pellet during density centrifugation. Hence, we sought to determine the degree of low-level blood contamination in the cellular and molecular fractions of the LN sample.

From the cellular perspective, we specifically examined the abundance of cells across each cluster contributed by blood or LN samples (Figure 2). Encouragingly, the Vδ2^+^ γδT cell, MAIT cell, CD14^+^ monocyte, and CD16^+^ monocyte clusters predominantly consisted of cells from the blood (Figure 2A), consistent with their reported paucity within SLOs (18, 19). Further supporting the notion that blood contamination was a minor concern, T_FH_ cells, all B cell populations, pDCs, and cDC1s were all over-represented in LN FNA-origin material (Figure 2A). To analyze this in a formal manner, we performed differential abundance analysis between blood and LN (see methods) (20). Blood origin cells were significantly enriched within the CD14^+^ monocyte and CD16^+^ monocyte clusters, as well as MAIT cell and Vδ2^+^ γδT cell clusters (Figure 2B and 2C). Conversely, neighborhoods consisting of switched (IgM^−^) and unswitched (IgM^+^) memory B cells, T_FH_ cells, and CD16^−^CD56^lo^ NK cells were enriched in LN.

**Figure 2.**
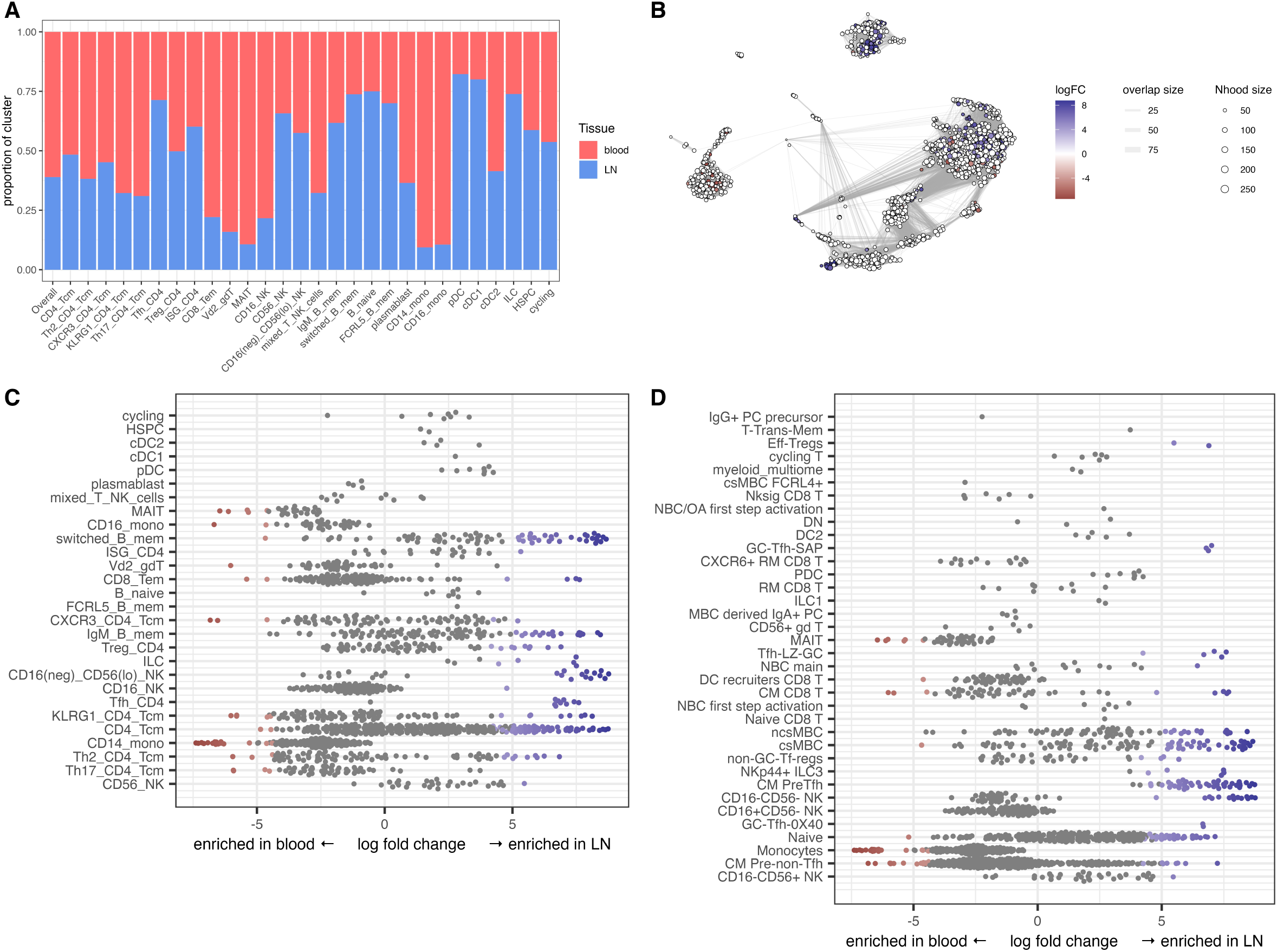
Differentially abundant populations between blood and lymph node. **(A)** Bar plot of the proportion of each cluster derived from LN or peripheral blood. **(B)** KNN weighted-network analysis to identify differentially abundant neighborhoods using the miloR package (see methods). **(C, D)** Relative abundance of neighborhoods based on cluster annotations (C) and on level 2 tonsil annotations (D). Neighborhoods enriched in cells from the blood are in red, neighborhoods enriched in cells from the LN are in blue, and no enrichment is in grey.

A particular strength of the *k*-nearest neighbor approach is the ability to examine changes in population abundance at a sub-cluster resolution (20). We utilized neighborhoods identified by this tool to annotate cells which did not form unique clusters but appeared potentially interesting in LN-specific populations (Figure 2D), based on Figure 1G. Using the tonsil atlas Level 2 annotations, non-class switched memory B cells (ncsMBC), class switched memory B cells (csMBC), and all identified T_FH_ populations (GC-Tfh-OX40, CM PreTfh, non-GC-Tf-regs, Tfh-LZ-GC, and GC-Tfh-SAP) were enriched in LN (Figure 2D). Furthermore, and consistent with the cluster-level analysis, MAIT cell and monocyte neighborhoods were enriched in blood whereas CD16^−^CD56^−^ NK cell-containing neighborhoods were enriched in LN (Figure 2D). Collectively, these data demonstrate LN FNA can efficiently recover tissue-specific populations including LN-specific innate immune cells.

### Differentially expressed genes and proteins between blood and lymph nodes

Next, we determined if the same cell clusters identified in blood and LN exhibited different transcriptional programs and expression of surface proteins. Nearly all clusters showed differentially expressed genes in blood or LN (Figure 3A), with expression changes most pronounced in the T and B cell clusters (Figure 3A). The high number of differentially expressed genes may reflect the presence of multiple distinct cell subsets with these clusters, based on the Azimuth annotations (Figure 1G) and neighborhood analysis (Figure 2D). The CD16^−^CD56^lo^ NK cell subset was the population with the greatest number of upregulated genes in both blood (246 genes up) and LNs (272 genes up; Supplemental Table 1). By contrast, the innate-like T cell populations (MAIT cells and Vδ2^+^ γδT cells) had relatively few differentially expressed genes, consistent with their highly conserved programs across tissues (21, 22). Innate immune populations were variable — monocytes, pDCs, and cDC1s differentially expressed few if any genes, while cDC2s (20 genes up in LN and 36 genes up in blood) and ILCs (17 genes up in LN and 14 genes up in blood) had greater differences between the two tissues (Figure 3A; Supplemental Table 1). Interestingly, these patterns did not correspond to differentially expressed proteins which showed no bias towards increased variability within the T or B cell clusters (Figure 3B). However, for nearly all populations, the LN cells showed significantly more upregulated proteins than blood (Figure 3B; Supplemental Table 2). There were a few genes (and corresponding proteins) that were enriched in nearly all populations in each tissue (Figure 3C): *CD69*, CD69 protein, *FOS*, and *JUN* were up in LN; *ITGA4* and CD49d protein in blood.

**Figure 3.**
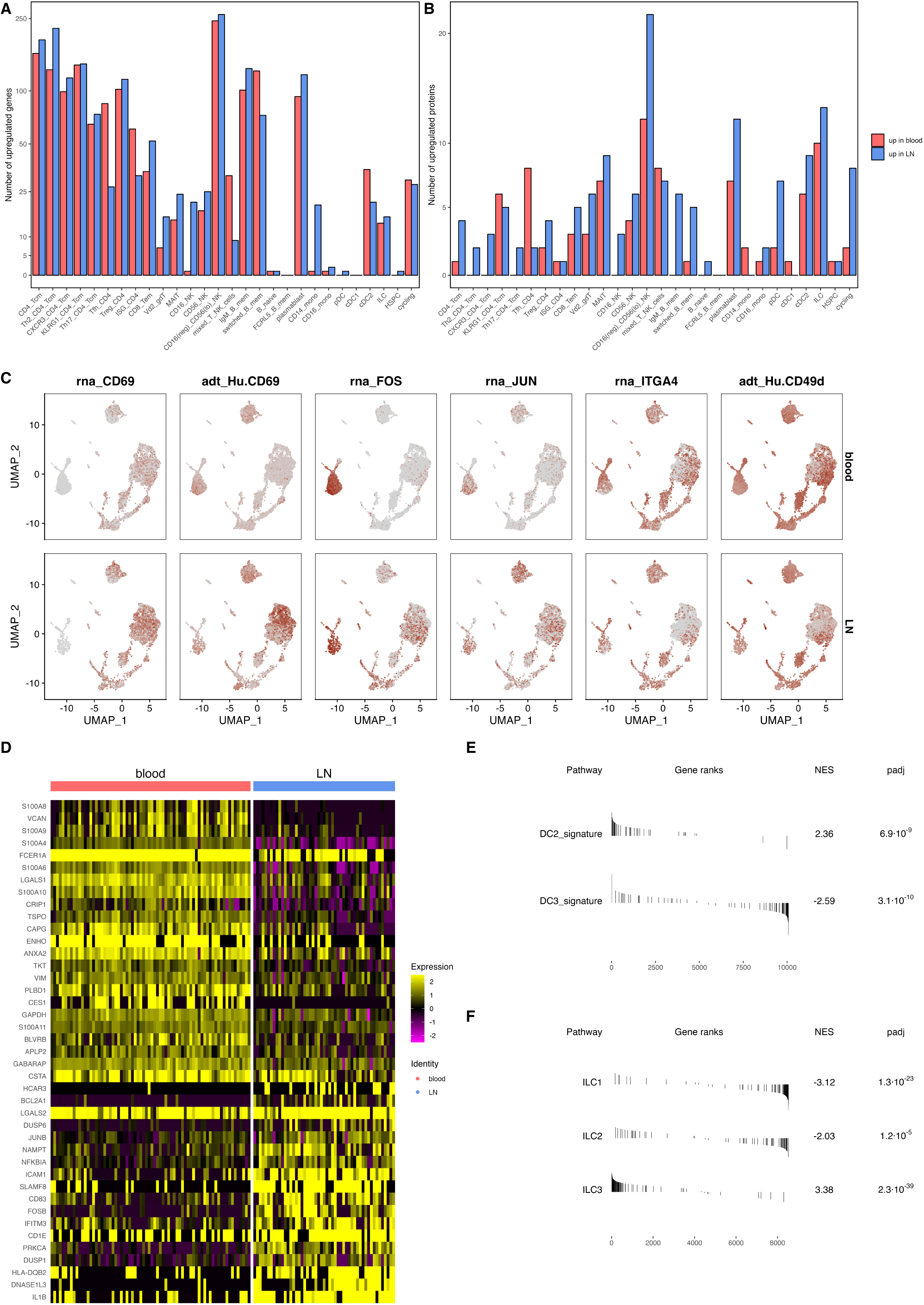
Differential expression of genes and surface proteins in lymph node cells. **(A)** Number of differentially expressed genes by cluster. **(B)** Number of differentially expressed surface proteins by cluster. **(C)** Overlay of selected genes (*CD69, FOS, JUN, ITGA4*) and proteins (Hu.CD69, Hu.CD49d) onto the UMAP, which is split by tissue of origin (top, blood; bottom, LN). **(D)** Differentially expressed genes on cDC2s from blood versus LN cells. **(E)** GSEA enrichment results comparing DC2 and DC3 signatures from (23) in lymph node versus blood cDC2s. **(F)** GSEA enrichment results comparing human ILC1, ILC2, and ILC3 signatures from (54) in lymph node versus blood ILCs. Positive NES (normalized enrichment score) values indicate enrichment in LN-origin cells and negative NES values indicate enrichment in blood-origin cells.

When studying DC populations critical for immune response initiation, we identified *FOS* as the only significantly differentially expressed gene between pDCs in blood and LN (up in LN; Figure 3D), and the nine significantly differentially proteins displayed gradations of expression between blood and LN (Supplemental Figure 3A). In contrast, the 56 differentially expressed genes and 15 proteins in the cDC2 cluster displayed a strong separation between blood and LN (Figure 3D, Supplemental Figure 3B). These genes broadly aligned with the differences between DC2 (e.g., *HLA-DQB2, CD1E, ARL4C*) and DC3 (e.g., *S100A8, VAN, S100A9*) subpopulations previously identified in human blood (23), which represent two states of CD1c^+^ DCs (“cDC2”) with lower or higher inflammatory signatures, respectively. Using GSEA analysis, the DC2 signature was confirmed to be enriched within LN (NES = 2.35, *P_adj_* = 4.4e^−9^), while the DC3 signature was enriched in blood (NES = −2.62, *P_adj_* = 9.3e^−11^; Figure 3E). These data suggest cDC2s exhibit a LN-specific profile, in contrast to cDC1 and pDCs that are similar across blood and lymphoid tissue.

ILCs consist of multiple transcriptionally distinct subsets with unique functions (24). We identified distinct ILC subsets between the two anatomic sites (Supplemental Figure 3B and 3C), with analysis showing a strong ILC3 signature within LN (NES = 3.38, *P_adj_* = 3.8e^−39^) and, conversely, strong signatures of ILC1s (NES = −3.22, *P_adj_* = 3.3e^−25^) and ILC2s (NES = −2.07, *P_adj_* = 3.4e^−5^) in the blood (Figure 3F). Thus, LN ILCs appear to be a distinct population from those found in circulation.

### LN-derived cells show greater predicted cell-cell interactions

One of our underlying hypotheses was that tissue sampling would reveal unique insights not available by studying blood. To test this, we utilized our single-cell sequencing data to examine predicted cell-cell interactions between cells within the blood or LN. CellPhoneDB (25, 26) predicted an overall greater number of cell-cell interactions between cells in the LN compared with blood (Figure 4A and 4B). In both tissues, most predicted interactions were between monocytes and NK cells. Compared with blood, LN T and B cell clusters had many more predicted interactions.

**Figure 4.**
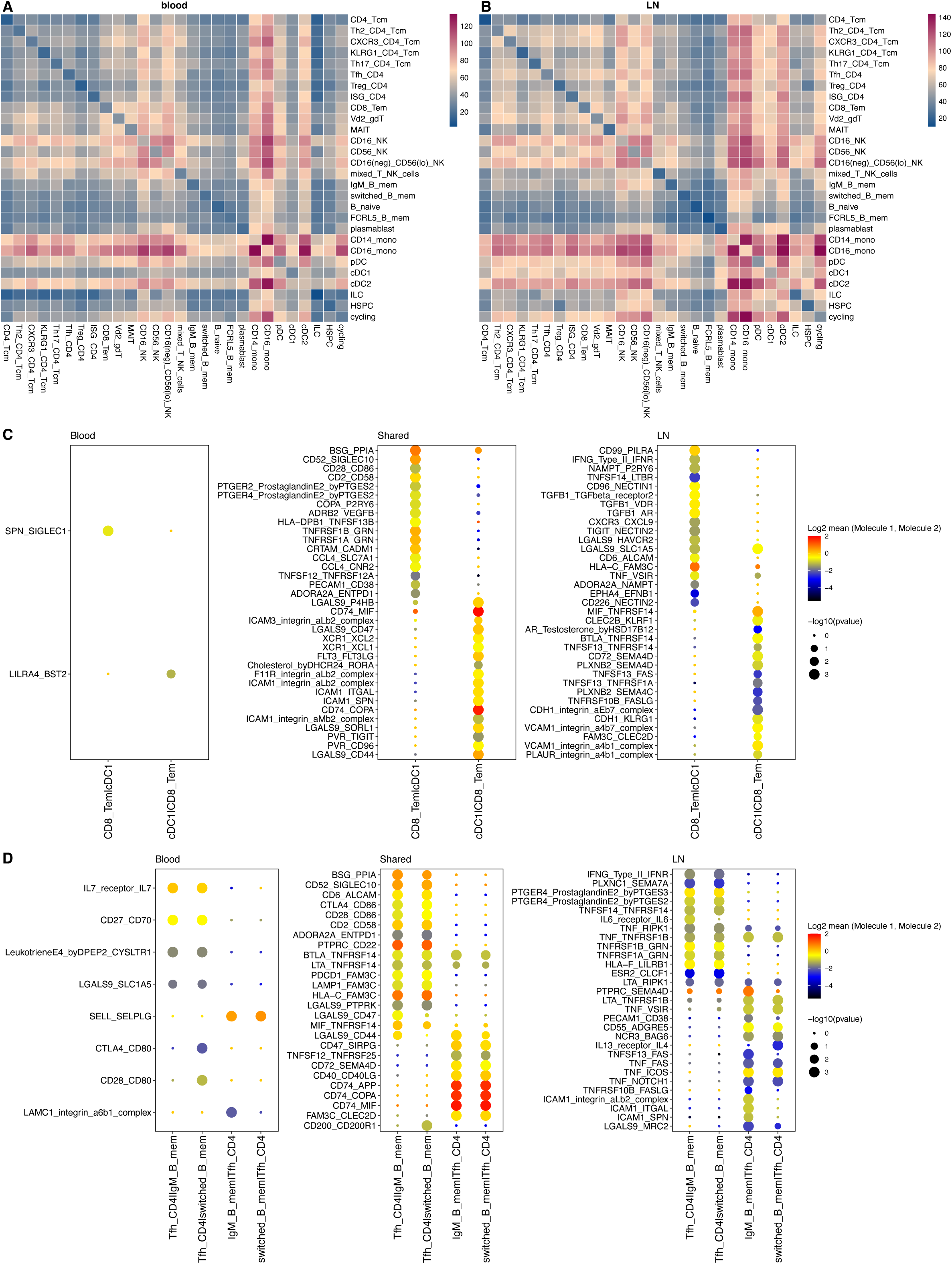
Unique predicted cell-cell interactions within the lymph node. CellPhoneDB was used to predict cell-cell interactions (see methods). **(A, B)** Total number of predicted significant interactions between each pair of cell types in blood (A) or LNs (B). **(C)** Predicted significant interactions between cDC1s and CD8^+^ T_EM_ cells based on tissue origin (blood, LN, or both tissues (“shared”)). **(D)** Predicted significant interactions between CD4^+^ T_FH_ cells and either IgM^+^ or Switched memory B cells based on tissue origin (blood, LN, or both tissues (“shared”)). Receptor-ligand pairs (y-axis) are written (receptor)_(ligand) and interacting cells (x-axis) are written (receptor-expressing cell) | (ligand-expressing cell).

To examine this more systematically, we examined predicted cell-cell interactions for populations with known biologic importance. For example, cDC1 cells have the unique ability to cross-present antigen, and thus play a critical role in the priming of CD8^+^ T cells (27). Accordingly, we examined predicted interactions between cDC1 cells and CD8^+^ T_EM_ cells and stratified these based on whether they were generated from our data on blood cells, LN cells, or from either source (Figure 4C). Thirty-five interactions were predicted regardless of which tissue source was used, including the traditional “signal 2” pathway of CD86-CD28 and the interaction of XCL1/XCL2 with XCR1 (the canonical marker of cDC1 cells (28)). Thirty-five additional interactions were predicted only by examining LN cells: these included IFNγ signaling via its receptor and CXCL9 interactions with CXCR3, a critical chemokine axis for CD8^+^ T cells (29). By contrast, only two interactions were uniquely predicted based on data from the blood.

Similarly, with regards to key GC-based T_FH_–B cell interactions, we identified a greater number of predicted pathways in the LN relative to blood (29 versus 6, with 26 additional pathways overlapping between tissues; Figure 4D). The canonical CD40L-CD40 signaling pathway (30), was predicted for the interaction of T_FH_ cells with both IgM^+^ and switched memory B cells, regardless of tissue origin. In contrast, IFNγ produced by T_FH_ cells was a uniquely predicted as a signaling pathway to both IgM^+^ and switched memory B cells in the LN and not in the blood. Differential interactions within a tissue were also predicted: IL-6 signaling was uniquely associated with signaling between T_FH_ cells and IgM^+^ memory B cells. While IL-4 produced by T_FH_ cells was uniquely associated with signaling to switched memory B cells, consistent with its established importance in class-switch recombination (31). Thus, these data validate and extend the importance of directly examining lymphoid tissue when considering pathways of interest for immune cell interactions.

### Lymph node aspirate supernatants are a source of unique proteins

Having validated the ability of cells collected by LN FNA to provide unique insights into SLO biology, we sought to determine if the aspirate supernatants could contribute orthogonal biologically pertinent information. To this end, we performed highly multiplexed proteomic analysis using the Olink Inflammation I and Neurology I assays using LN supernatants or blood. As two FNA passes (needles) were performed per participant, and each needle was flushed four times with PBS to facilitate cell recovery, we first sought to determine whether this impacted on protein recovery. The highest protein quantities (NPX) were recovered from the first flush of either needle, with particularly good recovery from the first wash of the second needle (LN needle 2; Figure 5A). Consistent with minimal blood contamination observed at the cellular level (Figure 2 and 3) and our previous observation of log-fold lower IgG in LN aspirates (7), proteomic analysis on capillary blood processed in the same manner as the FNA supernatant consistently recovered less material than LN needle 2 (Figure 5B). Additionally, an equivalent volume of serum also resulted in lower protein recovery (Figure 5B).

**Figure 5.**
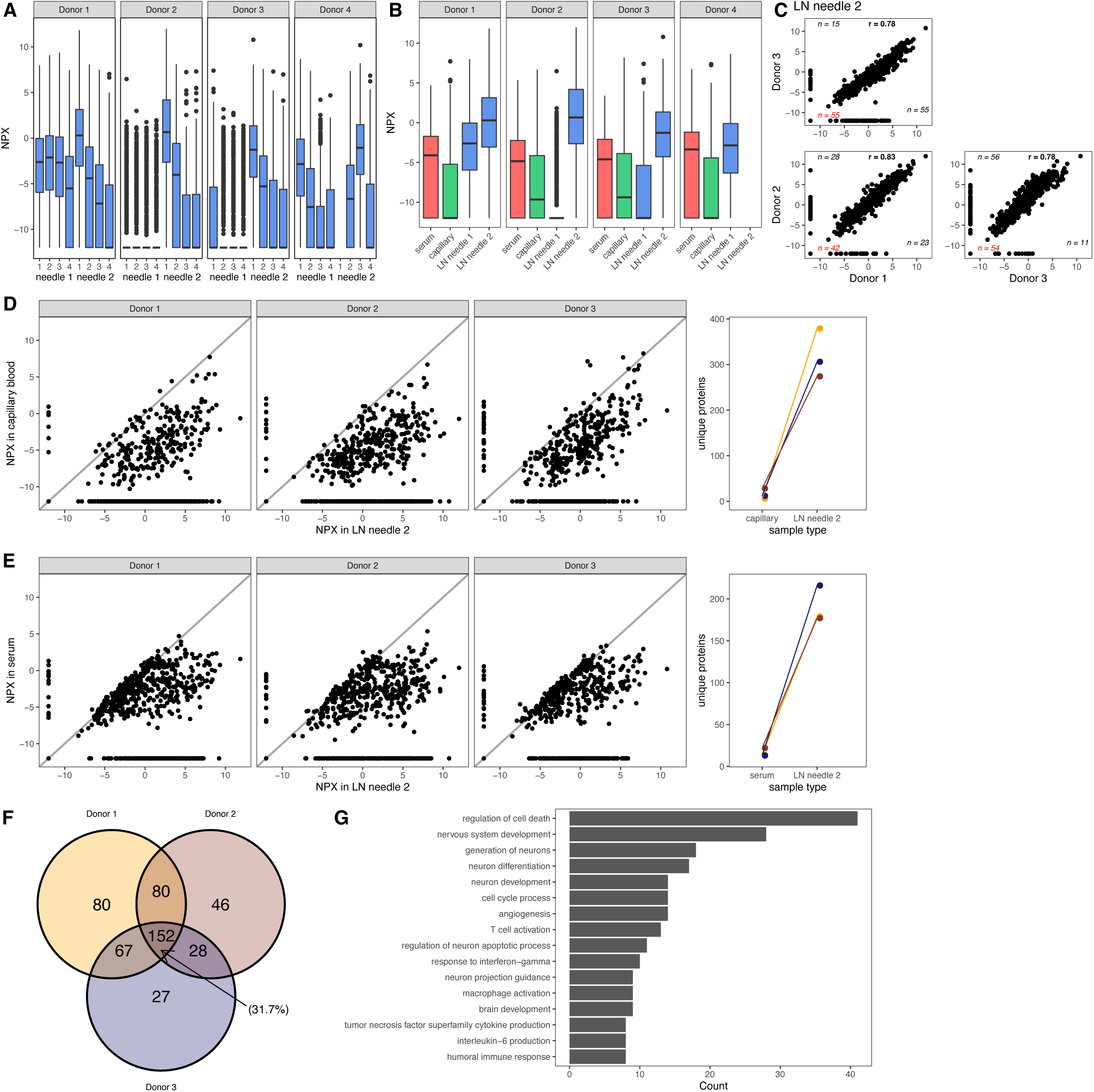
The FNA supernatant contains lymph node-specific proteins. Supernatant from LN FNA, matched serum, and matched cell-free capillary blood was subjected to proteomic analysis for 722 analytes (see methods). **(A)** Relative abundance of each protein after the first or second FNA needle, and each wash of the needle. Needle 2 wash 1 for Donor 4 was not analyzed. **(B)** Comparison of relative protein abundance in matched serum (diluted 1:100), capillary blood, LN FNA needle 1 (wash 1), and LN FNA needle 2 (wash 1). (A,B) Boxes indicate median, 25th percentile, and 75^th^ percentile. Whiskers are the IQR and dots represent outliers. **(C)** Correlation of relative protein abundance between LN FNA needle 2 (wash 1) for each donor. Number in black on each axis is the number of proteins not detected in the respective sample. Number in red in the bottom left is number of proteins not detected in either sample. Number in bold and black is Pearson correlation coefficient. **(D, E)** Left: correlation plots between matched LN FNA needle 2 (wash 1) and capillary blood (D) or serum (diluted 1:100) (E). Grey line is a line of identity. Right: number of proteins specifically detected in each matched sample are shown. Each color represents a donor. **(F)** Venn diagram of overlap of proteins specifically detected in each of the LN needle 2 (wash 1) samples compared to matched capillary blood. Number in parentheses indicates fraction of proteins found in all three samples. **(G)** GO pathway analysis of the 152 proteins specifically found in all three LN needle 2 (wash 1) samples compared to matched capillary blood. Selected pathways are shown. Count is number of proteins from the dataset contributing to each pathway.

Between donors, there was a high concordance of the recovered proteins in LN needle 2 and their concentrations (Figure 5C). There was a subset of proteins detected in blood but not in the FNA supernatants (Figure 5C, red text). The proteins differentially detected between participants showed overall lower abundances than the proteins readily detected in both samples of the pairwise comparison (*P* < 3.9e^−5^; Figure 5C). In contrast, the correlation of protein concentration and detection of specific proteins was much more variable for LN needle 1 (first wash of needle 1; Supplemental Figure 4A). Inter-sample correlation analysis of capillary blood (Supplemental Figure 4B) and diluted serum (Supplemental Figure 4C) showed highly concordant abundance patterns. However, there were a much larger number of proteins not detected in either of these samples as compared to the LN aspirates.

Next, we asked if lymphatic proteins draining into the node can be detected by FNA by comparing the proteins detected in the LN needle 2 and matched capillary blood. On average there were 300 proteins (of 722 total analytes) specifically detected in the LN aspirates (Figure 5D). Comparison of LN needle 2 and matched serum showed a slightly greater overlap, but still on average approximately 175 proteins were specifically detected by LN FNA (Figure 5E). LN needle 1 again proved less useful (Supplemental Figure 4D and E). From a comparison of the proteins specifically found in the LN aspirates (needle 2), 152 proteins were found in all 3 samples (31.7%; Figure 5F; Supplemental Table 3). A comparison of specifically detected proteins in all three LN aspirates relative to serum also found 31% overlap, consisting of 93 proteins (Supplemental Figure 4F).

Several of the proteins specifically detected in the LN aspirates of all three participants (Figure 5F) include TREM2, ADAM22, NTF3, and NMNAT1 (Supplemental Table 3), all of which have well-documented roles in neuronal and microglial function (32–35). As the cervical LNs sampled are thought to drain the dural sinuses (4), we postulated that detection of these proteins in LN aspirates would suggest sampling of lymphatic-associated material. To examine this more formally, ontology analysis was performed on the proteins specifically detected in the LN supernatant. In addition to enrichment for expected terms around immune function (e.g., T cell activation or humoral immune response), several terms associated with neuronal development, differentiation, and survival were enriched (Figure 5G). Many of the same terms were also enriched relative to serum (Supplemental Figure 4G). Thus, examination of the supernatant from LN aspirates is a fruitful means of identifying proteins specifically associated with the LN, whether they are produced locally or traffic from the lymph.

### Integration of proteomic and transcriptomic analysis

Finally, we sought to integrate proteomic and single-cell transcriptomic analyses towards LN specific receptor-ligand pathways. The list of 152 proteins specifically identified in LN (Figure 5F) was subset to a list of 96 proteins robustly detected within the transcriptomic data (Figure 6A). Expression levels of the genes were then visualized per cluster using only cells from the LN. Some genes such as *JUN, RABGAP1L*, and *IL16* had broad expression across the clusters (Figure 6A). Enrichment of *CLEC4C* was detected only in pDCs (Figure 6A), consistent with its use as a canonical marker of this population (36). Other interesting protein-producer cell pairings included CD200, which was strongly associated with T_FH_ cells and CCL3 which was strongly associated with the CD16^−^CD56^lo^ NK cell subset (Figure 6A).

**Figure 6.**
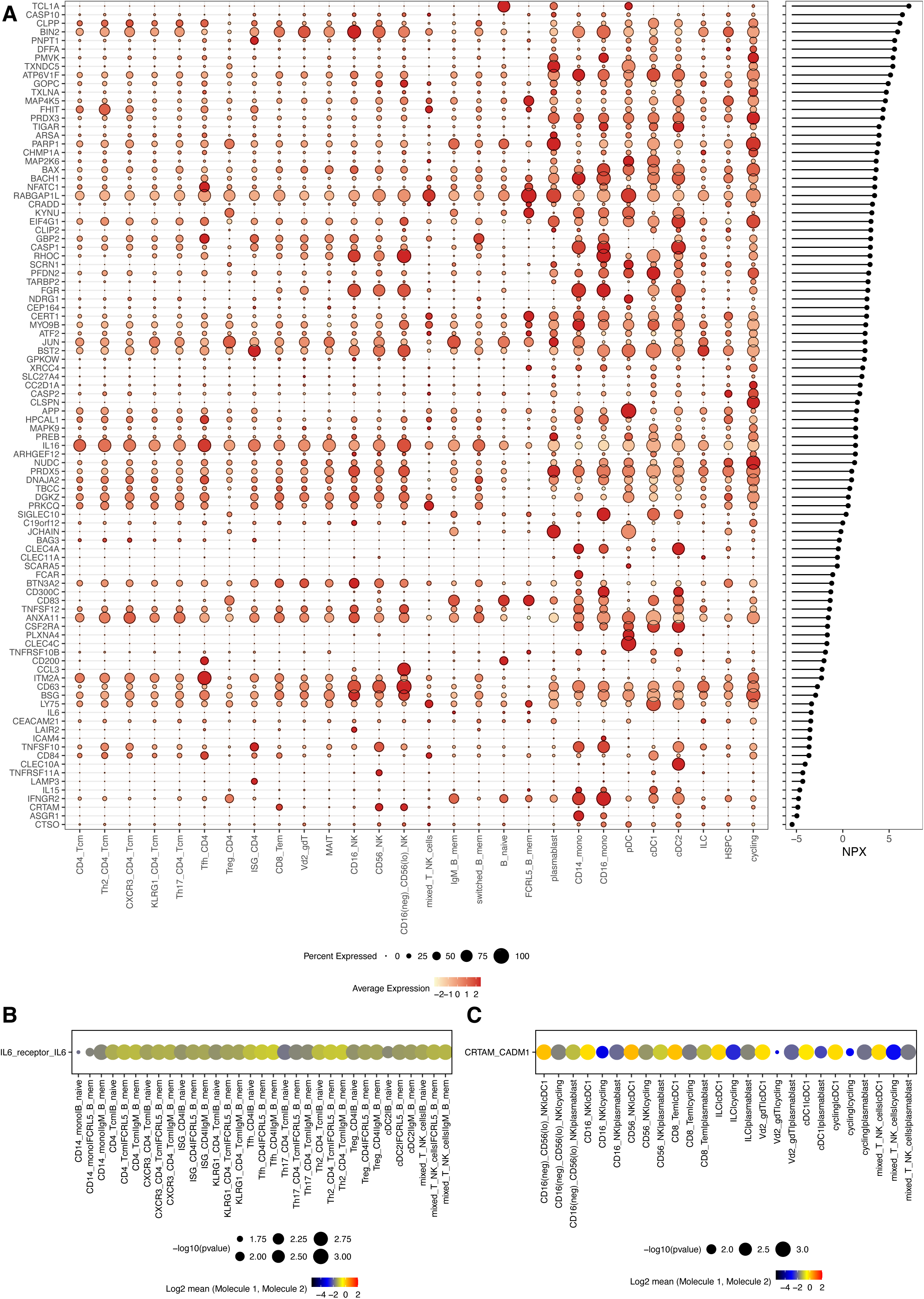
Proteomics can inform interpretation of single-cell transcriptomic analysis. **(A)** Dot plot expression of selected genes from LN cells corresponding to LN-specific proteins. Right plot depicts relative abundance of each protein. **(B, C)** CellPhoneDB predicted interactions of LN cells via the IL-6–IL-6 receptor signaling pathway (B) or the CADM1–CRTAM signaling pathway (C). Receptor-ligand pairs (y-axis) are written (receptor)_(ligand) and interacting cells (x-axis_ are written (receptor-expressing cell) | (ligand-expressing cell).

An alternative use for the proteomic data is to have it guide analysis of cell-cell interactions. For example, IL-6 was specifically detected in the LN aspirates (Supplemental Table 3), and examination of predicted cell-cell interactions using CellPhoneDB revealed all B cell clusters except switched memory B cells as possible sources of IL-6 with a potential signaling to essentially all CD4^+^ T cell clusters as well as CD14^+^ monocytes and cDC2s (Figure 6B). CRTAM was another protein specifically detected in supernatants of LN aspirates (Supplemental Table 3), which has a role in activation of CD8^+^ T cells and NK cells (37). Consistent with this, *CRTAM* was mostly highly expressed by CD8^+^ T cells and NK cells (Figure 6A). Its ligand (*CADM1*) was expressed by cDC1s, plasmablasts, and cycling cells (Figure 6C) – which appear to be cycling GC cells (Figure 1G) – and suggest a possible mechanism of NK-mediated regulation of GC function (38). In sum, merging these modalities can provide insights into specific cellular sources of LN proteins of interest or can inform on signaling pathways of interest through protein-guided validation.

## DISCUSSION

In this project, we sought to provide proof-of-concept of the utility of LN FNA to examine immune parameters beyond the main prior immunologic area of focus, GC B cell and T_FH_ cell biology, and into innate immune cells relevant to priming. We performed unbiased scRNA-seq on recovered cells and high-dimensional proteomics on FNA supernatants. Dendritic cell and other innate immune subsets were detected in sufficient numbers for downstream analysis. Predicted cell-cell interaction analysis highlighted many pathways only predicted by examining LN material, thus reinforcing the importance of studying this tissue. Proteomic analysis also revealed proteins otherwise undetectable by sampling peripheral blood, and these could be further used to cross-validate putative signaling pathways of interest.

A technically important question surrounding fine needle aspiration is to what extent blood contamination contributes to the recovered populations, given the vascularized nature of the target tissue alongside the sampling process. In contrast to studies of GC B cells or T_FH_ cells, this question is of more concern when examining cell types that have analogs in the circulation. Differential distribution of cell populations between blood and tissues has been the primary validator of the tissue-specific nature of the recovered sample. Prior work examining LN FNA purity has used very low abundance of granulocytes (11) or monocytes (8) as evidence. We confirm these findings through low abundance of monocytes, and also demonstrate low frequencies of MAIT cells and Vδ2^+^ γδT cells (18, 19). In addition, we identified an over-abundance of CD16^−^CD56^lo^ NK cells, which have been reported in SLOs but are rare in blood (17, 39). Our detection of specific proteins only within the FNA supernatant further supports the specific nature of this sampling technique. Thus, we are confident that innate immune populations recovered by LN FNA are of genuine LN origin.

Differential gene expression analysis of DC subsets revealed few differentially expressed genes and proteins for cDC1 and pDC populations in the LN compared to blood. Prior comparison of human DC phenotype and morphology between blood and LN DCs reported similarities between these tissues (40). scRNA-seq of blood DCs has revealed greater heterogeneity, especially within cDC2s and pDCs (23). At a clustering level, we could not identify populations beyond the historical XCR1^+^ cDC1, CD1c^+^ cDC2, and BDCA2^+^ pDC subsets. However, deeper investigation of the cDC2 cluster revealed distinct transcriptional programs suggesting a more “inflammatory” signature in blood origin cells and a “non-inflammatory” signature in LN origin cells. Thus, CD1c^+^ DCs (cDC2 subset by classical naming) in the LN and blood appear to be quite different populations. The relative enrichment of “non-inflammatory” signature CD1c^+^ DCs within the LN aligns with data from a recent tonsil atlas (17). That these co-clustered in our data may reflect a limitation of our sample size, given the relative scarcity of these cells (discussed below).

Despite detecting no differentially expressed genes between blood and LN origin cDC1s, a large number of cell-cell interactions were predicted only when examining LN-derived cells. This was primarily driven by the large number of differentially expressed genes on the second cell population (e.g., CD8^+^ T cells) in the interacting pair. These data would suggest that these lymphoid populations change phenotype upon entry into the lymph node. Presumably in the context of an ongoing immune response a very different conclusion would be drawn due to activation of the innate immune cells by PAMPs/DAMPs – future studies will be needed to investigate this hypothesis.

One of the goals of the current study was to investigate orthogonal uses for LN FNA beyond cellular analysis to aid in understanding of immune processes within the lymph node. Lymph contains a proteome distinct from peripheral blood (41, 42). Detection of proteins in the LN FNA supernatant that were not detectable from capillary blood or serum supports the conclusion that we are sampling lymph node tissue and/or lymph-associated proteins, as opposed to blood contaminants introduced by the procedure. There has been interest, particularly in cancer, in proteomic analysis of FNA supernatants – from possible malignant tissue or metastases containing LNs – to screen for the presence of cancer cells (43–45). Such studies focus on detecting cellular abundance (i.e., presence of cancer cells). Our detection of CLEC4C and its association with abundance of pDCs supports the use of proteomic analysis in this way for immune cell populations. Another unique way to merge cell- and protein-level data, is the ability to use proteomics to investigate active signaling pathways. We have previously detected CXCL13 within the LN, which was correlated with autoantibody production (6). Our ready detection of hundreds of proteins within the LN FNA supernatant suggests this is an approach with wide utility.

Detection of proteins in the supernatant more strongly associated with CNS tissue, such as TREM2, ADAM22, NTF3, and NMNAT1, suggests possible detection of lymph-associated proteins. However, some of these proteins are reported as detectable in lymphoid tissue (Human Protein Atlas; proteinatlas.org). This highlights a limitation of the current study: there was not sampling of a second lymph node site not associated with CNS drainage. Therefore, future studies with multiple anatomically separated sampling sites will be required to fully address this question. A second limitation of the current study is the relatively small sample size (number of donors and therefore number of cells). During ongoing immune reactions, lymph node cellularity increases substantially (46), which increases cell yields by FNA. There are multiple factors around the technique that impact on cell yields (11). Our participants did not have acute infection or ongoing neuroimmune disease, we only performed two FNA passes, and aspiration was not performed during the procedure. These factors all likely contributed to the number of cells recovered. Such factors should be carefully considered when designing an experimental plan, especially if studying rare cell populations (e.g., DCs).

In summary, we demonstrate the utility of LN FNA combined with omics approaches to study immune processes beyond the GC reaction. Such analysis of innate immune populations, and their interaction with adaptive immune subsets reveals biology not otherwise predicable based on blood sampling. These analyses can be augmented through use of LN FNA supernatants as a source of cell-free material to study pathways of interest. Finally, our dataset provides a useful reference for the cell populations, secreted proteins, and their abundances that can be reliably recovered by LN FNA, which provides a key benchmark when designing future studies using this approach. Detection of proteins associated with neuronal function is potentially compatible with detection of lymphatic drainage. Overall, these data provide a framework for future studies in the context of disease or other immune perturbations.

## METHODS

### Research ethics

Four donors (3 male, 1 female; 24 to 38 years of age) were consented for collection of lymph node material by fine needle aspiration (LN FNA) and matched blood (REC: 16/YH/0013). Informed written consent was received from all participants. All work was performed in accordance with relevant ethical regulation and was performed in compliance with the principles of the Declaration of Helsinki (2008).

### Lymph node sample collection and processing

Lymph node fine needle aspiration was performed on either level Ia/Ib (Donor 4) or Va (Donors 1-3) lymph nodes under ultrasound guidance as previously described (6). Two FNA passes were performed on the same lymph node using an identical approach. A sterile 23G 0.60×25 mm blue hypodermic needle (BD Bioscience) was inserted into the lymph node and via a repetitive corkscrew motion material was collected into the needle barrel via capillary action. The needle was withdrawn, and 1 ml of ice-cold, sterile PBS was aspirated through the bevel into a syringe. The needle was removed, and the material was released into a 1.5 ml microtube (Axygen). A total of 4 flushes of the needle were performed into separate collection tubes. The flushing process was repeated a total of four times into separate collection tubes. The process was then repeated for the second FNA pass using a fresh needle. All samples were stored on ice until processing.

Samples (8 tubes per participant) were spun down at 400 g for 5 min in a benchtop micro-centrifuge. Supernatant was carefully removed, transferred to a fresh 1.5 ml microtube, and stored at -80 °C for future use. Cell pellets were resuspended in 1 ml of R10 media (RPMI-1640 + 10% FBS + 1% Penicillin/Streptomycin) and combined into a single 15 ml conical. An additional 1 ml of R10 was used to wash and collect residual material. Samples were centrifuged at 931 g for 10 min, 4 °C. Supernatant was discarded and cells were resuspended in 2 ml of R10 and passed through a 30-micron filter into a 5 ml polystyrene FACS tube. An underlay of 2 ml of lymphoprep (STEMCELL technologies) was added. Cells were centrifuged at 931 g for 30 minutes with no brake. The mononuclear cell (MNC) layer was collected and transferred to a fresh 15 ml conical, which was topped up with 13 ml of R10. Cells were centrifuged at 754 g for 8 min and supernatant was discarded. Cells were resuspended in 500 μl of R10 and counted. The appropriate number of cells were used fresh for downstream applications.

### Capillary blood sample collection and processing

Capillary blood was collected from the index finger using a 1.5 mm safety lancet (Greiner Bio-one). Blood was collected into a 1.5 ml microtube. Blood was then drawn into a 23G 0.60×25 mm blue hypodermic needle (BD Bioscience), same as used for FNA, until the needle was full. The contents of the needle were flushed with 1 ml of ice-cold PBS into a fresh 1.5 ml microtube. This was done to replicate the experimental procedure used to collect LN FNA material. The diluted capillary blood was spun down at 400 g for 5 min in a benchtop micro-centrifuge. Supernatant was carefully removed, transferred to a fresh 1.5 ml microtube, and stored at -80 °C for future use.

### Veinous blood sample collection and processing

20 ml of peripheral blood was collected by venipuncture into an EDTA tube (BD Bioscience) for plasma and PBMC collection and 5 ml into an SST tube (BD Bioscience) for serum collection. After allowing to stand, SST tubes were spun down at 2000 g for 10 min. Serum was collected and frozen at -80 °C for future use.

To isolate peripheral blood mononuclear cells (PBMCs), EDTA blood was layered over 15 ml lymphoprep and centrifuged at 931 g for 30 minutes with no brake. The PBMC-containing layer was collected, transferred to a fresh 50 ml conical and topped up to 40 ml with R10 media. Samples were centrifuged at 754 g for 8 minutes and supernatant was discarded. Cells were washed an additional time with 40 ml of R10 before being counted. The appropriate number of cells were used fresh for downstream applications.

### Cell staining and FACS

Lymph node MNCs or PBMCs (5×10^5^ cells) were added to a 96-well U-bottom plate. Cells were centrifuged at 1,126 g for 2 min and supernatant discarded. Cells were washed once with FACS buffer (PBS + 0.05% bovine serum albumin + 1 mM EDTA) and centrifuged at 1,126 g for 1 min. Supernatant was discarded and cells were stained for 10 min at 4 °C with 2.5 μl Human TruStain FcX block (BioLegend) and 22.5 μl FACS buffer. After staining, cells were centrifuged at 1,126 g for 1 min and supernatant was discarded. Cells were stained with a cocktail containing CD3-PE (clone: UCHT1; 1:100), IgD-PE-Dazzle594 (clone: IA6-2; 1:100), CD19-PE-Cy7 (clone: HIB19; 1:100), CD27-APC (clone: O323; 1:50), CD45-AF700 (clone: HI30; 1:100), CD45RA-BV785 (clone: HI100; 1:200). All antibodies were from BioLegend. Final staining volume was 25 μl and staining was performed for 10 min at 4 °C. TotalSeq-C Human Universal Cocktail V1.0 (BioLegend) was prepared per manufacturer’s instructions and added to the cells without washing. Staining was performed for an additional 30 min at 4 °C. After staining, cells were washed 3 times with FACS buffer, resuspended in 200 μl of FACS buffer and transferred to a 1.5ml microtube. DAPI was added at a final concentration of 3 nM and samples were stored at 4 °C until sorting.

Sorting was performed on a BD FACSAria III (BD Bioscience) using an 85-micron nozzle. Three populations were collected: non-T/non-B (live, CD45^+^CD3^−^CD19^−^), non-naïve T cells (live, CD45^+^CD3^+^ and NOT CD45RA^+^CD27^+^), and non-naïve B cells (live, CD45^+^CD19^+^ and NOT IgD^+^CD27^−^). This sorting scheme resulted in a minor contamination of naïve B cells, confirmed in the scRNA-seq data, but nearly 100% purity of the other populations (Supplemental Figure 1). Cells were sorted into R20 (RPMI-1640 + 20% FBS + 1% Penicillin/Streptomycin) and kept at 4 °C. Flow cytometry data was analyzed using FlowJo (v 10.8.1; Becton Dickinson & Company).

### Single-cell RNA-sequencing library preparation

The three sorted cell populations were combined into a single tube, with 4,000 cells (or the entire population if < 4,000 sorted cells) from each population. Samples were centrifuged at 400 g for 5 minutes, washed once with loading buffer (PBS + 0.04% bovine serum albumin), and resuspended at an appropriate volume in loading buffer for incorporation into the 10x Genomics Chromium Next GEM Single Cell 5’ v2 (Dual Index) workflow. One sample (∼12,000 cells) was loaded per channel of the Chromium Controller. Gene expression libraries and cell surface protein (“ADT”) libraries were prepared according to manufacturer’s instructions. Sequencing was performed at the Oxford Genomics Centre (University of Oxford) on a Novaseq 6000 per manufacturer’s instructions. Illumina’s bcl2fastq was used to generate FASTQ files.

### Proteomic analysis

Lymph node FNA supernatant, capillary blood supernatant, and serum (pre-diluted 1:100 in PBS to approximate the volume of blood drawn by aspiration needle) were run using the Olink Explore 384 Inflammation I and Neurology I panels per manufacturer’s instructions (performed at the Oxford Genomics Centre). Proteins that were not detected in any samples were excluded from downstream analysis. Duplicate proteins (TNF, IL6, CXCL8) were removed from the dataset by keeping data only from the Neurology I panel. Proteins with an NPX value > −12 in the first sample and ≤ −12 in the second sample were defined as unique to the first sample. A series of t-tests (per donor pair) was performed to examine differences in NPX levels between proteins concordantly and discordantly detected across samples. The most conservative *P* value is reported. GO term analysis was performed using clusterProfiler (v 4.6.2)(47, 48) using the BP (Biological Process) ontogeny. Enriched sets were trimmed to include only those with > 5% of GO term list found in our query data. Particular GO terms of interest were manually selected from this list for visualization.

### Data QC and clean-up

Sequencing reads were mapped to the GRCh38-2020-A reference genome using CellRanger multi v7.0.1 and a custom marker list was provided for ADT mapping. Median reads per cell was between 36,000 and 262,000 for gene expression data and 3,300 and 20,000 for ADT data. Data was imported into Seurat (v 4.3.0) (16) and low-quality cells (UMI count < 900, RNA count < 1,300, or percent mitochondrial reads > 6.5%) were removed. Doublets were flagged and removed (except from the cycling cell cluster) using scDblFinder (v 1.12.0) (49). Immunoglobulin and T cell receptor gene segments, identified from the IMGT database (imgt.org), were removed from the gene expression object, except for the following genes: *IGHM, IGHG1, IGHG2, IGHG3, IGHG4, IGHD, IGHE, IGHA1, TRAV1-2, TRAV24, TRDV1, TRDV2, TRDV3*. Gene expression data were log normalized, variable features identified, and scaled using default Seurat parameters. The optimal number of PCs (45 dimensions) was determined by a combination of an elbow plot, and manual inspection of PCs. ADT protein data was normalized using the CLR method (margin = 2) and scaled. Clusering was performed using Seurat’s graph-based approach (Louvain algorithm). Harmony (v 0.1.1) (50) was used to integrate data across donors.

### Cluster annotation

Cluster annotation was performed using a combination of expert knowledge and the Azimuth web tool (azimuth.hubmapconsortium.org) (16). PBMC (16) and human tonsil (17) datasets were used as references. An optimal clustering resolution of 1.2 was selected with input from Clustree (v 0.5.0). A multimodal weighted nearest neighbor integration approach (16) was investigated, but was not felt to improve cluster annotation, so was not pursued (not shown).

### Differential abundance analysis

Differential abundance testing was performed using miloR (v 1.7.1) (20). Specified parameters were k = 30, d = 30, prop = 0.1. The design was ∼ tissue. plotDAbeeswarm was used without removing mixed populations.

### Differential gene expression

Differential gene expression and differential protein expression analysis was performed using MAST (51) using the Seurat FindMarkers function on a per-cluster basis comparing blood and lymph node as the two conditions, with donor as a latent variable. Significant DEGs were defined as padj < 0.05. Differentially expressed genes and proteins were visualized using FeaturePlot or DoHeatmap.

### Gene Set Enrichment Analysis

Gene Set Enrichment Analysis (GSEA) (52) was performed using the fgsea package (53). Gene signatures for DC2 and DC3 subsets are from the DEGs listed in (23). The gene signature for ILC subsets was generated by taking the uniquely expressed genes for each subset from (54) and selecting only significantly upregulated genes within each cell subset. TCR gene segments, and ribosomal genes were removed, and the list was trimmed to 150 genes for ILC3s (ILC1 and ILC2 signatures already contained < 150 genes). Data were visualized using plotGseaTable.

### Cell-cell interactions

Predicted cell-cell interactions were defined using CellPhoneDB (v 3.10) (25, 55) with database version 4.0.0. Blood and lymph node data were separately extracted from the Seurat object and run using cellphonedb method statistical_analysis --counts_data hgnc_symbol --threshold 0.1. Data visualization was performed by adapting the plotting functions included in the CellPhoneDB package (https://github.com/ventolab/CellphoneDB).

### Data availability

All sequencing data is deposited in the GEO database (accession number XXXX). Analysis pipelines are available upon reasonable request.

## ACKNOWLEDGMENTS

We thank the donors for their participation. We thank Helen Ferry for assistance with cell sorting. Analysis was performed using the MRC WIMM Centre for Computational Biology computing cluster. NMP is supported by a Wellcome Career Development Award [227217/Z/23/Z] and a Goodger & Schorstein Scholarship. AAD is funded by a NIHR Clinical Lectureship and Academy of Medical Sciences Starter Grant for Clinical Lecturers (SGL027\1016). PK is supported by a Wellcome Senior Fellowship [222426/Z/21/Z]. SRI is supported by the Wellcome [104079/Z/14/Z], the Medical Research Council (MR/V007173/1), BMA Research Grants – Vera Down grant (2013), Margaret Temple (2017), Epilepsy Research UK (P1201), the Fulbright UK-US commission (MS Society research award), and by the National Institute for Health Research (NIHR) Oxford Biomedical Research Centre. The views expressed are those of the author(s) and not necessarily those of the NHS, the NIHR or the Department of Health.

## SUPPLEMENTAL MATERIAL

**Supplemental Figure 1.**

Sort scheme for isolation of cells for single-cell sequencing.

**Supplemental Figure 2.**

**(A)** Dot plot showing normalized protein expression of the indicated proteins across each of the clusters. Characters after the dash (-) indicate the antibody clone for selected antibodies. **(B, C)** Annotation scores from Azimuth for blood (B) and LN (C) using the Level 2 PBMC, Level 1 Tonsil, and Level 2 Tonsil annotations.

**Supplemental Figure 3.**

**(A, B)** Differentially expressed proteins in pDCs (A) and cDC2s (B) between blood and LN cells. **(C, D)** Differentially expressed genes (C) and proteins (D) in the ILC cluster between blood and LN cells.

**Supplemental Figure 4.**

**(A-C)** Correlation of relative protein abundance between LN FNA needle 1 (wash 1) (A), capillary blood (B), and serum (diluted 1:100) (C) for each donor. Number in black on each axis is the number of proteins not detected in the respective sample. Number in red in the bottom left is number of proteins not detected in either sample. Number in bold and black is Pearson correlation coefficient. **(D, E)** Number of proteins specifically detected in each matched sample of (D) capillary blood versus LN needle 1 (wash 1) or (E) serum (diluted 1:100) versus LN needle 1 (wash 1). Each color represents a donor. **(F)** Venn diagram of overlap of proteins specifically detected in each of the LN needle 2 (wash 1) samples compared to matched serum (diluted 1:100). Number in parentheses indicates percent of proteins found in all three samples. **(G)** GO pathway analysis (ontogeny = biological process) of the 92 proteins specifically found in all three LN needle 2 (wash 1) samples compared to matched serum (diluted 1:100). Selected pathways are shown. Count is number of proteins from the dataset contributing to each pathway.

**Supplemental Table 1.** Differentially expressed genes between blood and LN, separated by cluster. Fold change > 0 indicates increased expression in LN and fold change < 0 indicates increased expression in the blood.

**Supplemental Table 2.** Differentially expressed proteins between blood and LN, separated by cluster. Fold change > 0 indicates increased expression in LN and fold change < 0 indicates increased expression in the blood.

**Supplemental Table 3.** Proteins specifically detected in supernatants of LN aspirates from all three participants relative to matched capillary blood.

