## Supplementary figures and images for "Fine needle aspiration of human lymph nodes reveals cell populations and soluble interactors pivotal to immunological priming"

### Supplemental Figure 1

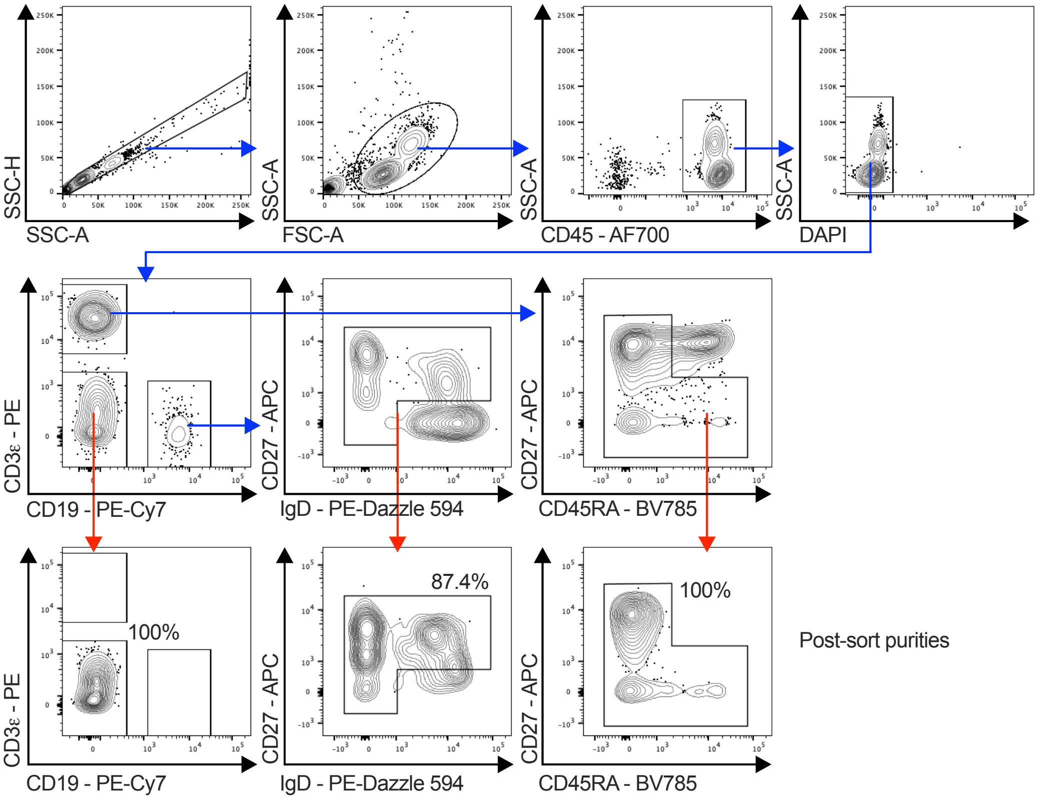

### Supplemental Figure 2

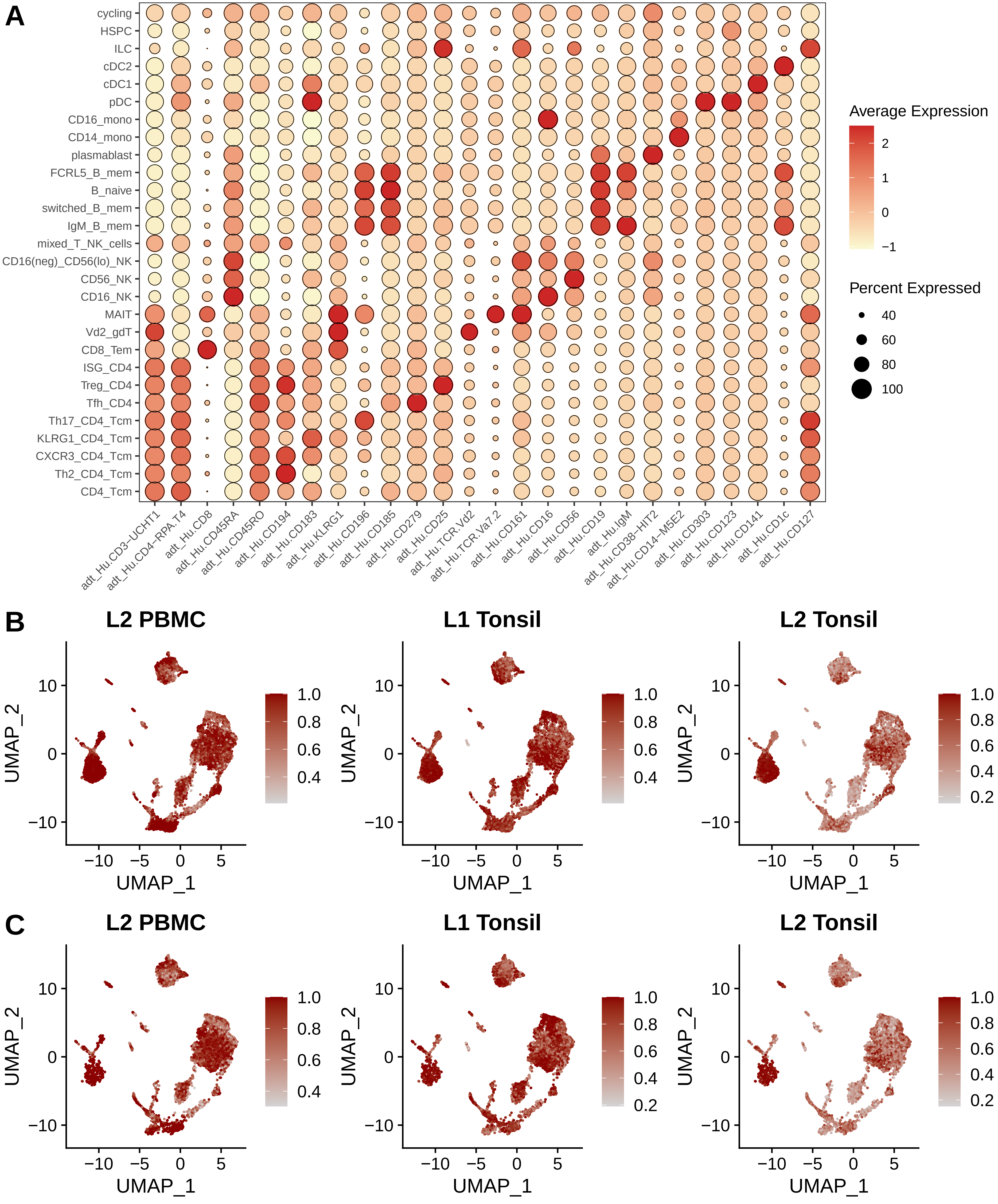

### Supplemental Figure 3

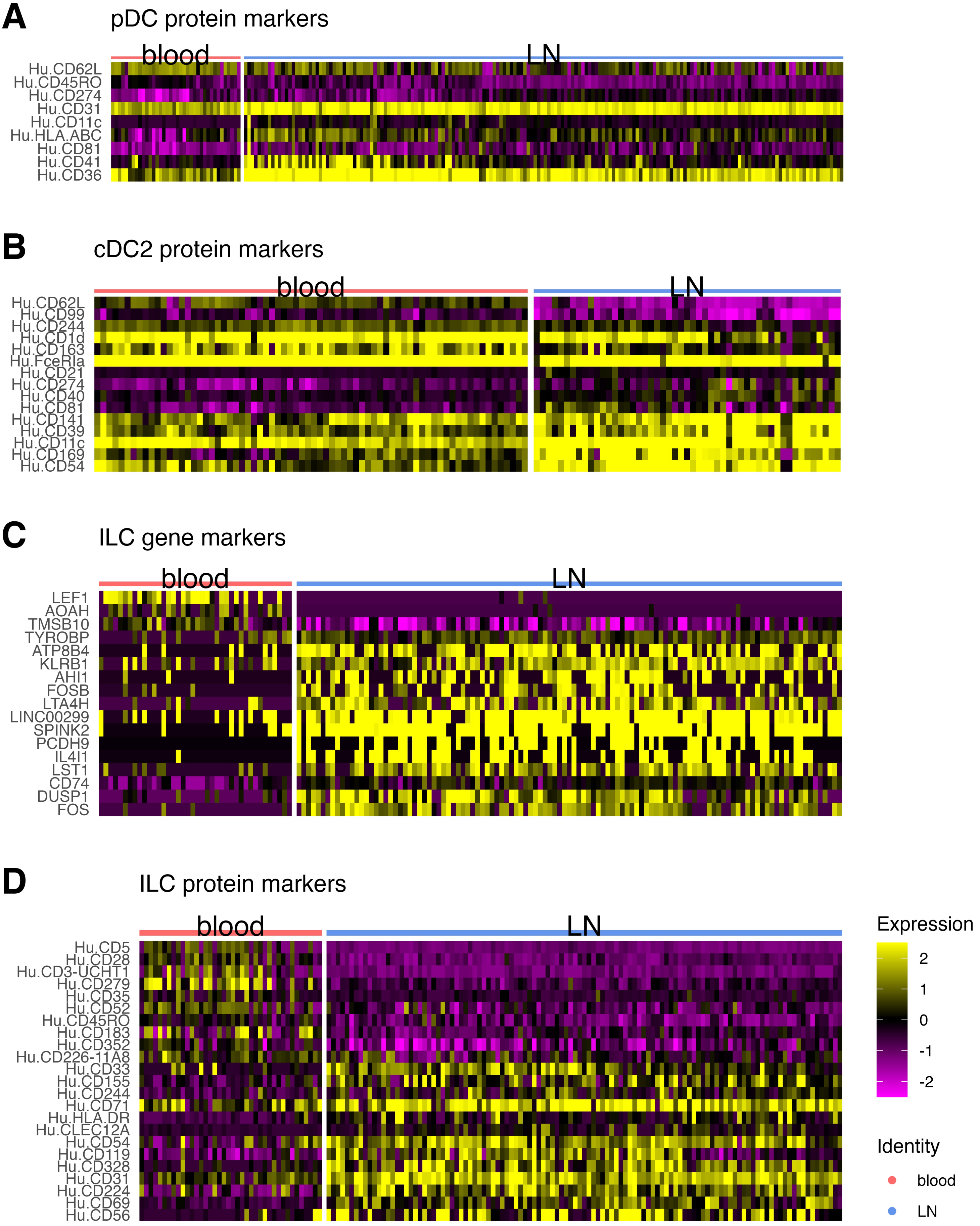

### Supplemental Figure 4

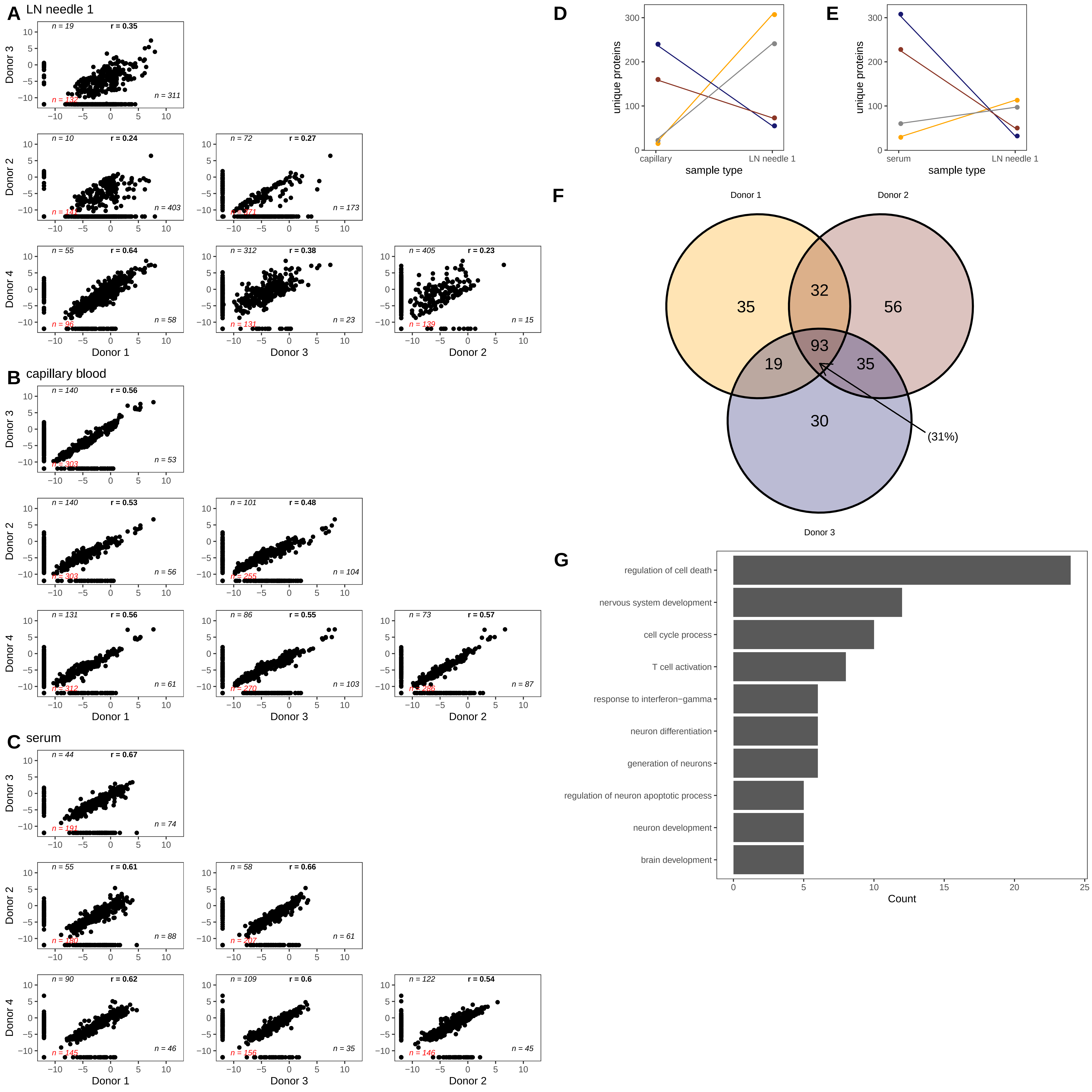
